# Gene Set Enrichment Analysis (GSEA) of Upregulated Genes in Cocaine Addiction Reveals miRNAs as Potential Therapeutic Agents

**DOI:** 10.1101/303362

**Authors:** Ishtiaque Ahammad

## Abstract

Cocaine addiction is a global health problem that causes substantial damage to the health of addicted individuals around the world. Dopamine synthesizing neurons in the brain play a vital role in the addiction to cocaine. But the underlying molecular mechanisms that help cocaine exert its addictive effect have not been very well understood. Bioinformatics can be a useful tool in the attempt to broaden our understanding in this area. In the present study, Gene Set Enrichment Analysis (GSEA) was carried out on the upregulated genes from a dataset of Dopamine synthesizing neurons of post-mortem human brain of cocaine addicts. As a result of this analysis, 3 miRNAs have been identified as having significant influence on transcription of the upregulated genes. These 3 miRNAs hold therapeutic potential for the treatment of cocaine addiction.

## Introduction

Cocaine addiction is a public health problem that spans the whole world. Formation of ‘molecular memory’ through various epigenetic mechanisms leading to neural gene expression is a means of drug addiction [1]. There are many types of cells in the brain which are associated with drug abuse but the most prominent of them dopamine synthesizing neurons of the ventral midbrain. Despite being relatively small in number among the neural cellular population, dopamine synthesizing neurons are significant in mediating both the acute rewarding effects of drugs of abuse and the conditioned responses to cues associated with previous drug use [4].

One study has shown that the psychostimulant actions of cocaine come from its ability to promote the release of dopamine into the synaptic cleft via exocytosis of synaptic vesicles containing dopamine [2]. Another study has revealed that a α1 receptor mediates the increase in dopamine levels elicited by cocaine [3].

Due to the critical role played by dopamine synthesizing neurons in addiction, shedding more light on drug-induced molecular changes in these cells has become crucial yet our understanding of the nature of these changes still remains far from complete [5]. For gaining new insight into the pathophysiology of complex disorders such as drug addiction, postmortem human brain can be a unique resource despite the challenges associated with its use [6].

In this study, upregulated genes in DA neurons from postmortem human brain were identified to shed more light on the molecular mechanism, pathways and the key players involved with cocaine addiction. For doing this, gene expression dataset GSE54839 was used [7].

## Methods

### Identification of Upregulated Genes from GSE54839

From the NCBI website GEO datasets were searched using the term “cocaine AND differential gene expression” and reference series GSE54839 was analyzed with GEO2R.

For GEO2R analysis two groups termed “Cocaine addiction” and “Control” were defined. Thirty samples belonged to each groups. Using the GEOquery [8] and limma R [9] packages from the Bioconductor project, GEO2R analysis was performed [10]. Top 250 differentially expressed genes were found. P values were adjusted using the Benjamini & Hochberg (false discovery rate) method [11]. Log2 transformation to the data was applied. R script used to perform the calculation was obtained from the R script tab.

### Enrichment Analysis

Enrichment analysis of the Upregulated genes was carried out using
- ChEA2016 TFs [12]
- MiRTarBase 2017 [13]
- KEGG 2016 [14]

## Results and Discussion

### R script

~~~
# Version info: R 3.2.3, Biobase 2.30.0, GEOquery 2.40.0, limma 3.26.8
# R scripts generated Tue Apr 17 00:24:53 EDT 2018
~~~

~~~
################################################################
# Differential expression analysis with limma
library(Biobase)
library(GEOquery)
library(limma)
~~~

~~~
# load series and platform data from GEO
~~~

~~~
gset <- getGEO(“GSE54839”, GSEMatrix =TRUE, AnnotGPL=TRUE)
if (length(gset) > 1) idx <- grep(“GPL6947”, attr(gset, “names”)) else idx <- 1
gset <- gset[[idx]]
~~~

~~~
# make proper column names to match toptable
fvarLabels(gset) <- make.names(fvarLabels(gset))
~~~

~~~
# group names for all samples
gsms<-“111000111000111000111000111000111000111000111000111000111000”
sml <- c()
for (i in 1:nchar(gsms)) { sml[i] <- substr(gsms,i,i) }
~~~

~~~
# log2 transform
ex <- exprs(gset)
qx <- as.numeric(quantile(ex, c(0., 0.25, 0.5, 0.75, 0.99, 1.0), na.rm=T))
LogC <- (qx[5] > 100) | |
     (qx[6]-qx[1] > 50 && qx[2] > 0) ||
     (qx[2] > 0 && qx[2] < 1 && qx[4] > 1 && qx[4] < 2)
if (LogC) { ex[which(ex <= 0)] <- NaN
  exprs(gset) <- log2(ex) }
~~~

~~~
# set up the data and proceed with analysis
sml <- paste(“G”, sml, sep=“”) # set group names
fl <- as.factor(sml)
gset$description <- fl
design <- model.matrix(~ description + 0, gset)
colnames(design) <- levels(fl)
fit <- lmFit(gset, design)
cont.matrix <- makeContrasts(G1-G0, levels=design)
fit2 <- contrasts.fit(fit, cont.matrix)
fit2 <- eBayes(fit2, 0.01)
tT <- topTable(fit2, adjust=“fdr”, sort.by=“B”, number=250)
~~~

~~~
tT <- subset(tT, select=c(“ID”,“adj.P.Val”,“P.Value”,“t”,“B”,“logFC”,“Gene.symbol”,“Gene.title”))
write.table(tT, file=stdout(), row.names=F, sep=“\t”)
~~~

~~~
################################################################
# Boxplot for selected GEO samples
library(Biobase)
library(GEOquery)
~~~

~~~
# load series and platform data from GEO
~~~

~~~
gset <- getGEO(“GSE54839”, GSEMatrix =TRUE, getGPL=FALSE)
if (length(gset) > 1) idx <- grep(“GPL6947”, attr(gset, “names”)) else idx <- 1
gset <- gset[[idx]]
~~~

~~~
# group names for all samples in a series
gsms<-”111000111000111000111000111000111000111000111000111000111000”
sml <- c()
for (i in 1:nchar(gsms)) { sml[i] <- substr(gsms,i,i) }
sml <- paste(“G”, sml, sep=“”) set group names
~~~

~~~
# order samples by group
ex <- exprs(gset)[, order(sml)]
sml <- sml[order(sml)]
fl <- as.factor(sml)
labels <- c(“cocain”,“control”)
~~~

~~~
# set parameters and draw the plot
palette(c(“#dfeaf4”,“#f4dfdf”, “#AABBCC”))
dev.new(width=4+dim(gset)[[2]]/5, height=6)
par(mar=c(2+round(max(nchar(sampleNames(gset)))/2),4,2,1))
title <- paste (“GSE54839”, ‘/’, annotation(gset), “ selected samples”, sep =“)
boxplot(ex, boxwex=0.6, notch=T, main=title, outline=FALSE, las=2, col=fl)
legend(“topleft”, labels, fill=palette(), bty=“n”)
~~~

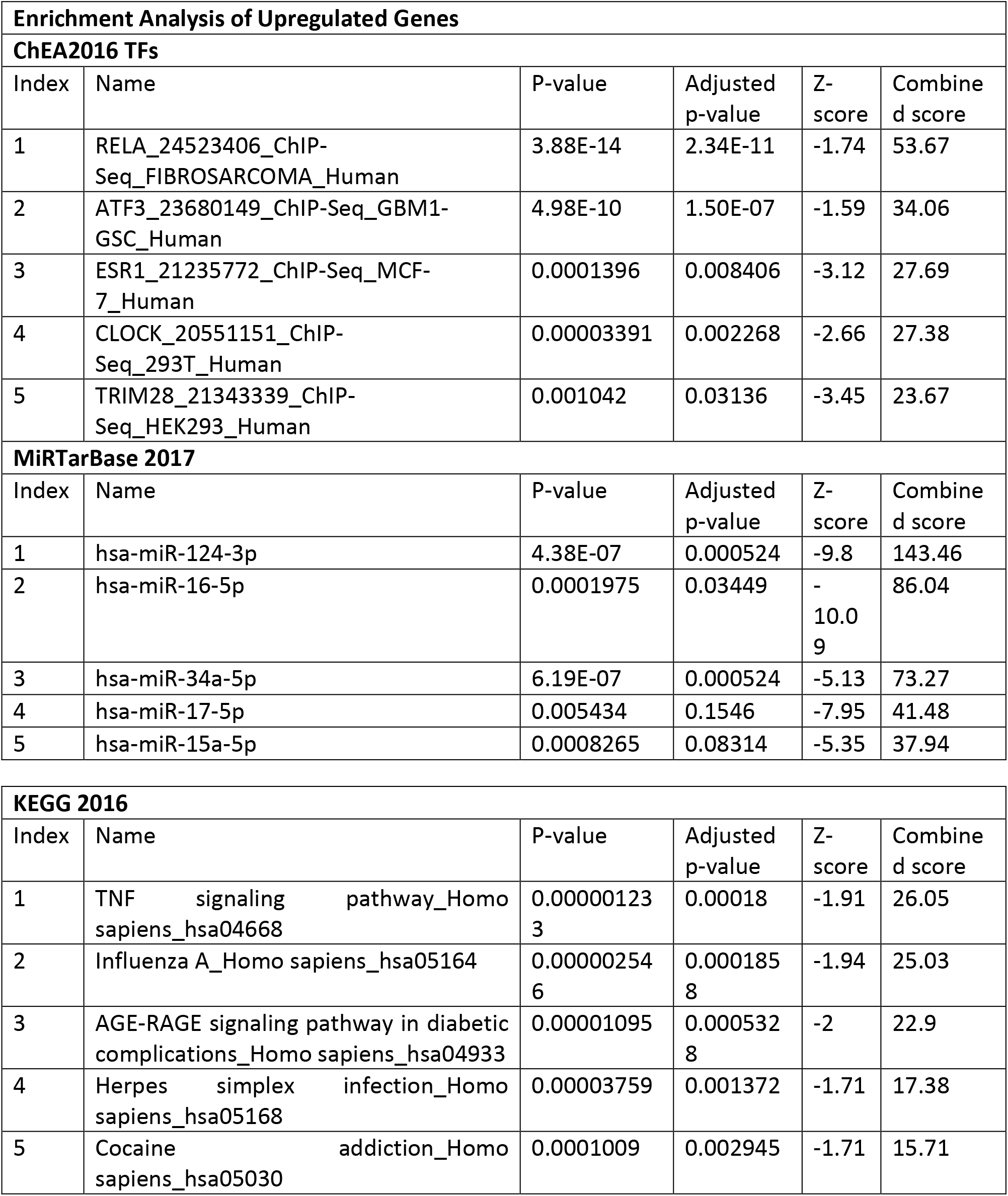

For the upregulated genes:

From the ChEA2016 TFs database, RELA was found to be the most significant transcription factor. Hsa-miR-124-3p, hsa-miR-16-5p and hsa-miR-34a-5p were identified as the top 3 most significant miRNAs from the MiRTarBase 2017. From the KEGG 2016 pathway analysis, TNF signaling pathway was found to be the most significant pathway mediated by the upregulated genes.

Precursor of the mature miRNA, hsa-miR-124-3p, namely the miR-124 is a small non-coding RNA molecule which has been fund in flies [15], nematode worms [14], mouse [13] and human [16]. Dicer enzyme processes the mature ~21 nucleotide mature miRNAs from hairpin precursor sequences. MiR-124 is the most abundant miRNA expressed in neuronal cells. The sequence for hsa-miR-124-3p is- 53 - uaaggcacgcggugaaugccaa - 74 [17]

Precursor of the mature miRNA, hsa-miR-16-5p namely the miR-16 family is vertebrate specific and its members have been predicted or discovered in a number of different vertebrate species. The sequence for hsa-miR-124-3p is- 14 - uagcagcacguaaauauuggcg - 35 [17]

Precursor of the mature miRNA, hsa-miR-34a-5p namely the miR-34 family gives rise to three major mature miRNAs. Members of the miR-34 family were discovered computationally at first [18] and verified experimentally later [19], [20]. The sequence for hsa-miR-34a-5p is- 22 - uggcagugucuuagcugguugu - 43 [17]

Role of miRNAs as important regulatory agents for gene expression is being considered as therapeutic means in various diseases. Unlike siRNAs, miRNA-targeted therapy is capable of influencing not only a single gene, but entire cellular pathways or processes. Mitigating the effects exerted by overexpression of malignant miRNAs is possible through the application of artificial antagonists such as oligonucleotides or other small molecules. It is also possible to supplement miRNAs through the use of synthetic oligonucleotides [21]. In the case of current study, the miRNAs which were found to influence the transcription of upregulated genes in cocaine addiction can be supplemented so that they can negatively regulate those genes and thus reduce the addictive effects of cocaine.

## Conclusion

From the Gene Set Enrichment Analysis, 3 miRNAs have been discovered to be related to the upregulated genes in cocaine addiction. Therefore it can be predicted that these 3 miRNAs hold therapeutic promise against cocaine addiction through silencing of the upregulated genes. Further studies *in vitro* and *in vivo* should be carried out in order to get more knowledge about the efficacies of these miRNAs in mitigating the effects of cocaine addiction.

